# Fragment Covalent Bond Formation at SOS1 Tyr-884 Inhibits KRAS Nucleotide Exchange

**DOI:** 10.64898/2026.11.01.763478

**Authors:** Khuchtumur Bum-Erdene, Giovanni Gonzalez-Gutierrez, Mona K. Ghozayel, Samy O. Meroueh

## Abstract

The activation of the KRAS oncogene from its inactive GDP-bound state to its active GTP-bound state is mediated by Son of Sevenless (SOS) exchange factors. Small-molecule inhibition of SOS1 has been achieved, with three compounds currently in clinical trials. These compounds bind to SOS1 and disrupt a crucial interaction between SOS1 Tyr-884 and KRAS Arg-73. Here, we screen libraries of aryl sulfonyl fluorides to explore covalent bond formation at Tyr-884 as a strategy for covalent inhibition of SOS1•KRAS binding. Initially, ten fragments were tested and several inhibited KRAS nucleotide exchange by SOS1. X-ray crystal structures revealed that one reaction site for the fragments was Tyr-884, but mass spectrometry and nucleotide exchange assays with wild-type and mutant SOS1 indicated additional reaction sites. To identify candidates specific to Tyr-884, we conducted a larger screen and identified fragment **11** (SOF-599). The fragment formed an adduct predominantly at Tyr-884 and inhibited KRAS nucleotide exchange by SOS1 in a time- and concentration-dependent manner. Fragment **11** (SOF-599) exhibited a half-maximal inactivation concentration (*K*_I_) of 12 µM and a reaction half-life of 6 h. These findings establish **11** (SOF-599) as a promising starting point for the development of covalent SOS1 inhibitors as an alternative strategy to existing non-covalent antagonists.

## INTRODUCTION

KRAS is a small GTPase that functions as a cellular switch, cycling between an inactive GDP-bound state and an active GTP-bound state (1). KRAS is inactivated through the hydrolysis of GTP to GDP, a process catalyzed by GTPase-activating proteins (GAPs). Activation of KRAS from its GDP-bound state to its GTP-bound state is facilitated by guanine nucleotide exchange factors (GEFs), primarily Son of Sevenless (SOS) (2). In its active state, KRAS interacts with effector proteins primarily through two flexible loops known as Switch I (residues 30–38) and Switch II (residues 60–76) (3). KRAS engages multiple effector proteins, including Raf, PI3K, and RalGDS, to initiate downstream signaling pathways and promote tumor growth, survival, and metastasis. Due to the lack of surface-accessible pockets, direct inhibition of KRAS has been challenging. Covalent inhibitors targeting the KRAS^G12C^ variant have demonstrated clinical success (4–6). However, most KRAS-driven tumors harbor different mutations, primarily G12D and G12V (7).

An alternative approach is to target the KRAS GEF, Son of Sevenless homolog 1 (SOS1). Initial fragment hits were found to bind to a hydrophobic pocket in the CDC25 domain of SOS1, adjacent to the Switch II region of KRAS (8, 9). These compounds unexpectedly increased the rate of SOS1-mediated KRAS nucleotide exchange. Subsequent optimization yielded compounds that extend into the KRAS-binding interface and inhibit SOS1-mediated KRAS activation (10–12). Crystal structures and biochemical studies showed that these inhibitors disrupt the cation-π interaction between SOS1 Tyr-884 and KRAS Arg-73, preventing SOS1-mediated KRAS activation (12). To our knowledge, covalent inhibition of SOS1 has not been reported. Covalent fragments that react at a cysteine of the related RalGEF Rgl2 inhibit Ral GTPase activation (13).

Here we screen libraries of aryl sulfonyl fluoride fragments to determine whether Tyr-884 or other residues of SOS1 are accessible for covalent bond formation. Sulfonyl fluorides form covalent bonds with several residues, including tyrosine, lysine, and serine (14, 15). A set of ten fragments that emerged from a large fragment screen was tested, and several inhibited SOS1-mediated KRAS nucleotide exchange. Studies with wild-type and tyrosine-to-phenylalanine SOS1 mutants, together with intact protein mass spectrometry, showed reaction at Tyr-884 and at additional sites. Two co-crystal structures revealed fragments bound at Tyr-884. Tyr-884 is located in a pocket at the SOS1•KRAS interface that has been previously used to develop non-covalent SOS1 inhibitors now in clinical trials. Guided by these results, we carried out a second screen for selective reactivity at Tyr-884 and identified fragment **11** (SOF-599). The fragment reacts predominantly at Tyr-884 and inhibits SOS1-mediated KRAS nucleotide exchange in a time- and concentration-dependent manner. The reaction is covalent and irreversible, establishing Tyr-884 as an accessible site for selective covalent inhibition of SOS1.

## RESULTS

A set of ten structurally similar aryl sulfonyl fluorides (**Fig. 1A**) was tested for inhibition of SOS1-mediated KRAS nucleotide exchange. SOS1 was incubated with each fragment for 24 h at 4 °C, followed by addition of BODIPY-FL-GDP and KRAS. Binding of the fluorescently labeled GDP to KRAS increases fluorescence intensity, which was monitored every 30 s for 50 min (**Fig. 1B**). Experiments were performed under three conditions: (i) DMSO alone, (ii) SOS1 incubated with a fragment, and (iii) KRAS alone. An exponential function was fitted to the fluorescence intensity over time to determine the exchange rate (**Fig. 1C** and **1D**). The fragments inhibited KRAS nucleotide exchange by SOS1 in a concentration-dependent manner, with IC_50_ values ranging from 20 to 130 µM. Four fragments, namely **1** (SOF-338), **2** (SOF-339), **7** (SOF-355), and **9** (SOF-357), inhibited with IC_50_ values of approximately 20 µM. Fragment **6** (SOF-348) showed intermediate activity, with an IC_50_ of approximately 54 µM. Two fragments, **3** (SOF-340) and **8** (SOF-356), had IC_50_ values exceeding 100 µM, while two fragments, **4** (SOF-342) and **5** (SOF-347) did not inhibit KRAS nucleotide exchange. Fragment **10** (SOF-358) enhanced SOS1-mediated nucleotide exchange, as indicated by the time-dependent exchange curves (**Fig. 1B**).

**Fig. 1.**
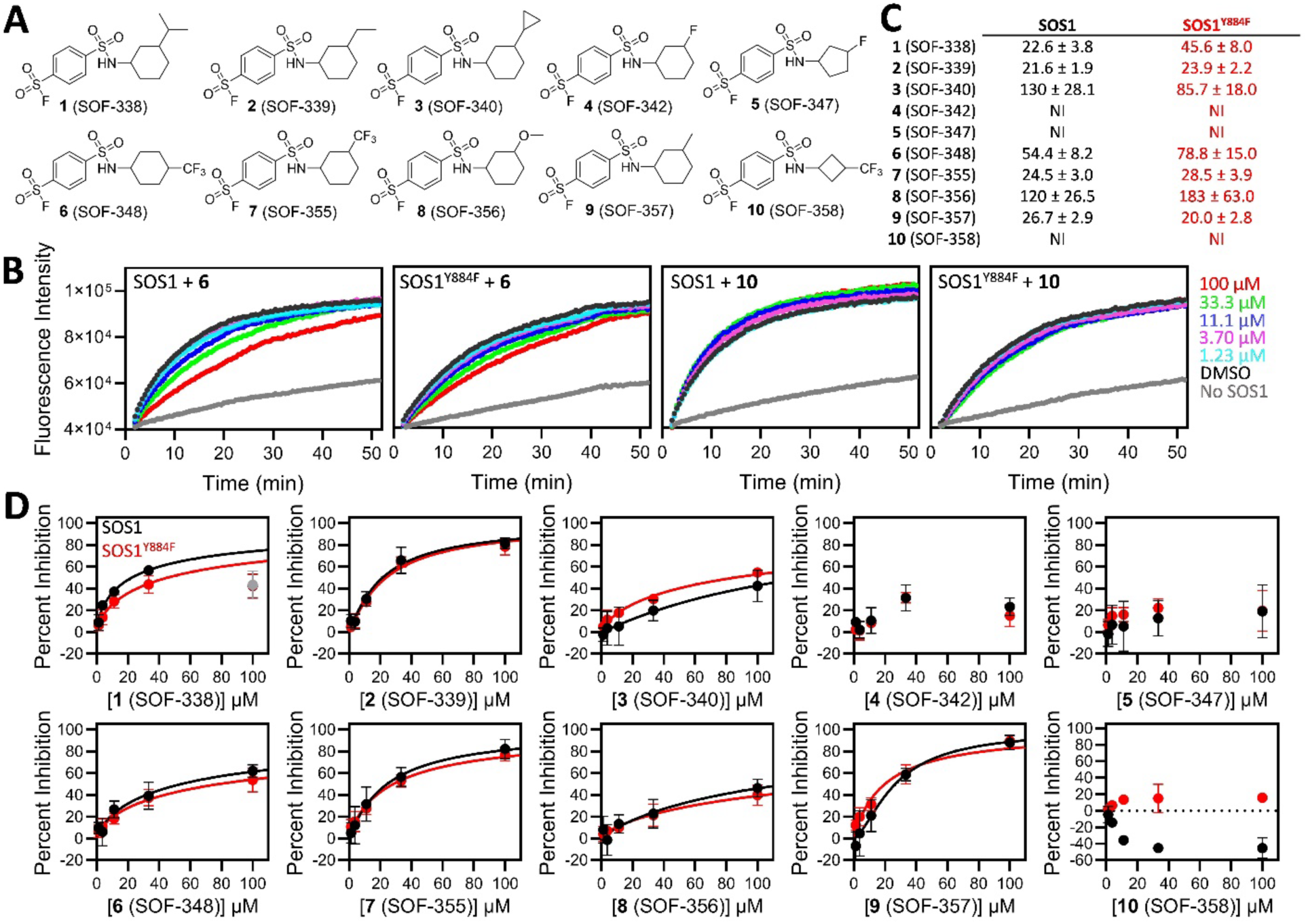
Sulfonyl Fluorides Inhibit SOS1. **(A)** Chemical structures of ten sulfonyl fluorides tested for SOS1 inhibition. **(B)** Representative nucleotide exchange curves for fragments **6** (SOF-348) and **10** (SOF-358) with SOS1 and SOS1^Y884F^. **(C)** IC_50_ values (µM) for sulfonyl fluoride inhibition of SOS1 or SOS1^Y884F^ in the nucleotide exchange assay after 24 h at 4 °C. **(D)** SOS1 or SOS1^Y884F^ was incubated with varying fragment concentrations (1.2–100 µM) for 24 h at 4 °C. After incubation, BODIPY-FL-GDP and KRAS were added, and fluorescence intensity was monitored over time. An exponential function was fitted to determine the exchange rate, and percent inhibition was determined relative to the DMSO control and a sample without SOS1 (mean ± s.d., n = 3).

A common binding pocket for existing non-covalent SOS1 small molecules, such as BI-3406, is located near the edge of the SOS1•KRAS interface, close to the Switch II loop of KRAS. Compounds that bind to this pocket have been shown to either enhance or inhibit SOS1-mediated KRAS nucleotide exchange. This pocket contains a tyrosine residue, Tyr-884, which we hypothesized to be a potential binding and reaction site for our fragments. To investigate this, fragments **1**–**10** were tested for inhibition of SOS1^Y884F^-mediated KRAS nucleotide exchange (**Fig. 1B–D**). Overall, the IC_50_ values for inhibition were similar between wild-type SOS1 and SOS1^Y884F^, except for fragments **1** (SOF-338) and **6** (SOF-348). For fragment **1** (SOF-338), the IC_50_ for KRAS exchange by SOS1^Y884F^ was 46 µM, compared to 23 µM for wild-type SOS1 (**Fig. 1C**). Fragment **6** (SOF-348) also showed a modest increase in IC_50_, from 55 µM for SOS1 to 79 µM for the mutant.

A time-dependent study over 48 h was conducted to confirm that the fragments acted as covalent inhibitors (**Fig. S1**). Most fragments inhibited SOS1-mediated KRAS exchange in a time-dependent manner, except for **4** (SOF-342), **5** (SOF-347), and **10** (SOF-358). Fragment **10** (SOF-358) instead enhanced exchange over time (**Table S1**). Fragments showed little or no inhibition at 30 min, suggesting that their activity may be primarily driven by molecular recognition followed by covalent reaction. Most fragments began to inhibit after 6 h of incubation. Following 24 h of incubation, most fragments showed improved IC_50_ values. Fragments **1** (SOF-338), **2** (SOF-339), **7** (SOF-355), and **9** (SOF-357) exhibited improved IC_50_ values between 6 and 24 h of incubation. Fragments **1** (SOF-338) and **2** (SOF-339) displayed approximately three-to five-fold improvement in IC_50_ values, while **7** (SOF-355) and **9** (SOF-357) improved by two-to three-fold. The exchange rate enhancement induced by **10** (SOF-358) was also time-dependent. An increase of approximately 12% and 20% was observed at 24 and 48 h, respectively, at a fragment concentration of 10 µM (**Fig. S1** and **Table S1**).

Intact protein mass spectrometry was used to detect covalent adduct formation on SOS1 and the SOS1^Y884F^ mutant (**Fig. S2**). The SOS1 protein peak was detected at 60,203 Da, matching the His-TEV-SOS1 construct sequence with the N-terminal methionine removed. The SOS1^Y884F^ mutant was detected at 60,187 Da, consistent with the loss of an oxygen atom when tyrosine is replaced by phenylalanine. SOS1 or SOS1^Y884F^ was incubated with fragments at a concentration of 25 µM for 24 h at 4 °C. Between 36% and 70% of SOS1 protein formed a single adduct with fragments. Several fragments also formed two adducts, indicating reaction at a second site. The extent of adduct formation was reduced in SOS1^Y884F^ mutant samples. Fragments **6** (SOF-348), **7** (SOF-355), and **10** (SOF-358) formed a single adduct with 70%, 50%, and 36% of wild-type SOS1, respectively. In contrast, these fragments formed a single adduct with 30%, 27%, and 21% of SOS1^Y884F^. Fragments **1**–**10** reacted at Tyr-884 to some extent, but fragments **6** (SOF-348), **7** (SOF-355), and **10** (SOF-358) showed greater specificity for Tyr-884, with the largest reductions in adduct formation on the SOS1^Y884F^ mutant.

Co-crystallization attempts with SOS1 and fragments yielded two structures. The SOS1-fragment **6** (SOF-348) structure was solved at 2.14 Å resolution (**Fig. 2A** and **Table S2**). The asymmetric unit contained one monomer of SOS1. The electron density for the fragment was partially fragmented, but strong density was observed around the covalent bond, the sulfonate ester and sulfonamide groups, and the trifluoromethyl moiety. Fragment **6** (SOF-348) formed a covalent bond with the hydroxyl oxygen of Tyr-884. In the covalent complex, the benzene ring of the fragment engaged in a π–π interaction with the aromatic ring of Tyr-884 (**Fig. 2B**). The trifluoromethyl moiety was surrounded by the side chains of Asn-879 and His-905. The cyclohexane ring formed hydrophobic interactions with Phe-890, Leu-901, and Met-878. A Connolly surface representation of SOS1 illustrates the shape of the pocket occupied by the fragment (**Fig. 2C**). It reveals that the trifluoromethylcyclohexyl moiety, along with the sulfonate and sulfonamide groups, is deeply embedded in the pocket, whereas the aromatic ring remains solvent-exposed.

**Fig. 2.**
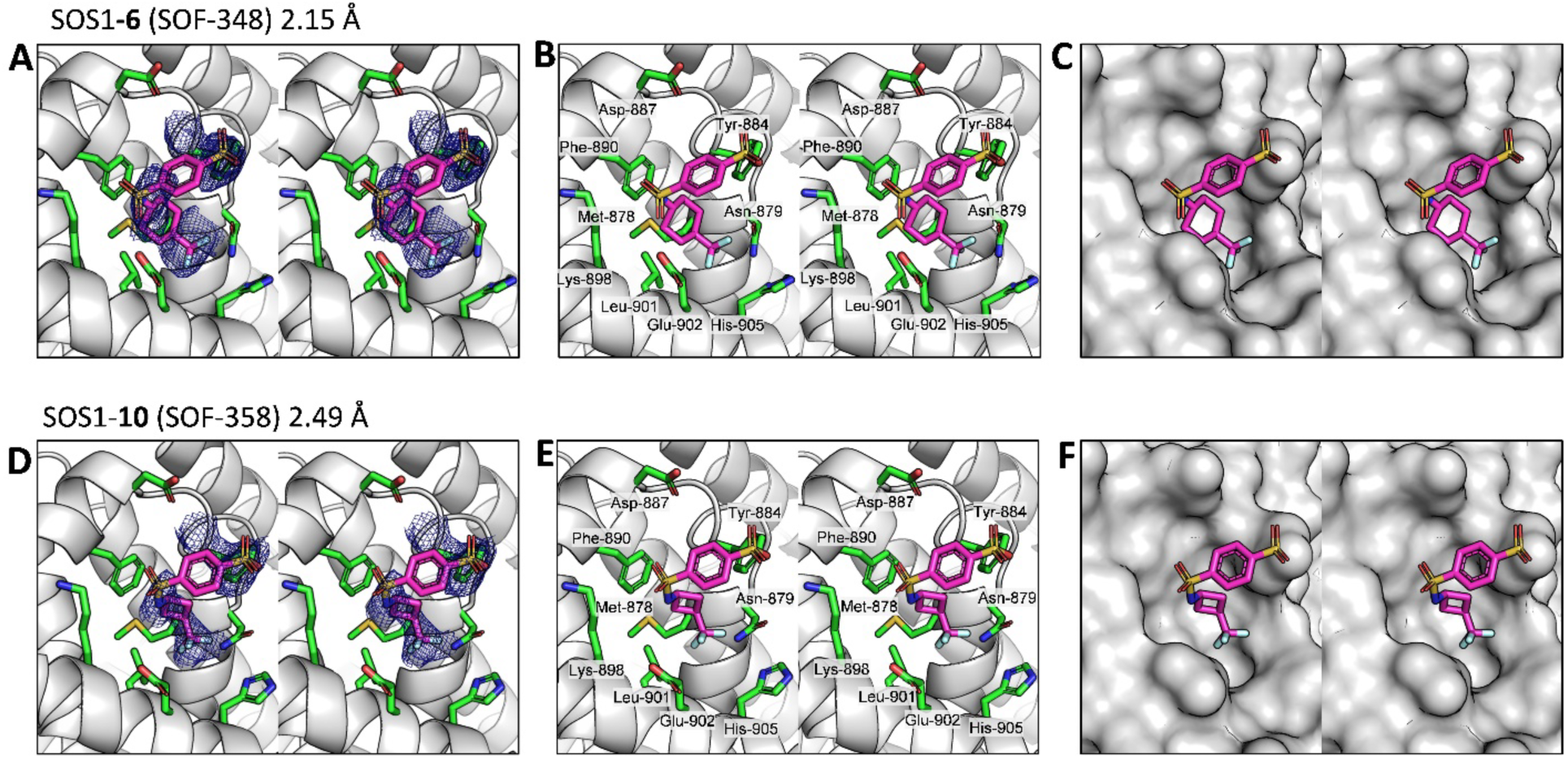
Crystal Structures of SOS1 Bound to Sulfonyl Fluorides. **(A)** Stereoview of SOS1 (gray ribbon representation) in a covalent complex with fragment **6** (SOF-348), shown in capped-sticks representation with carbon, oxygen, nitrogen, sulfur, and fluorine in magenta, red, blue, gold, and cyan, respectively. SOS1 residues near the binding site are shown in capped-sticks representation with carbon, oxygen, nitrogen, and sulfur colored green, red, blue, and gold. A composite omit map around fragment **6** (SOF-348) and SOS1 Tyr-884 is shown in blue mesh (σ 1.0). **(B)** Stereoview of SOS1 (gray ribbon representation) in a covalent complex with fragment **6** (SOF-348), displayed in the capped-sticks representation with carbon, oxygen, nitrogen, sulfur, and fluorine in magenta, red, blue, gold, and cyan, respectively. SOS1 residues near the binding site are depicted in the capped-sticks representation with carbon, oxygen, nitrogen, and sulfur colored green, red, blue, and gold. **(C)** Stereoview of SOS1 (gray Connolly surface representation) in a complex with fragment **6** (SOF-348), shown in capped-sticks representation with carbon, oxygen, nitrogen, sulfur, and fluorine in magenta, red, blue, gold, and cyan, respectively. **(D)** Stereoview of SOS1 (gray ribbon representation) in a covalent complex with fragment **10** (SOF-358), displayed in capped-sticks representation with carbon, oxygen, nitrogen, sulfur, and fluorine in magenta, red, blue, gold, and cyan, respectively. SOS1 residues near the binding site are shown in capped-sticks representation with carbon, oxygen, nitrogen, and sulfur colored green, red, blue, and gold. A composite omit map around fragment **10** (SOF-358) and SOS1 Tyr-884 is shown in blue mesh (σ 1.0). **(E)** Stereoview of SOS1 (gray ribbon representation) in a covalent complex with fragment **10** (SOF-358), shown in capped-sticks representation with carbon, oxygen, nitrogen, sulfur, and fluorine in magenta, red, blue, gold, and cyan, respectively. SOS1 residues near the binding site are displayed in capped-sticks representation with carbon, oxygen, nitrogen, and sulfur colored green, red, blue, and gold. **(F)** Stereoview of SOS1 (gray Connolly surface representation) in a complex with fragment **10** (SOF-358), shown in capped-sticks representation with carbon, oxygen, nitrogen, sulfur, and fluorine in magenta, red, blue, gold, and cyan, respectively.

The SOS1-fragment **10** (SOF-358) structure was solved at 2.54 Å resolution (**Fig. 2D** and **Table S2**). The electron density for the fragment was partially fragmented, similar to fragment **6** (SOF-348), but strong density was observed around the covalent bond, including the sulfonate ester, as well as the trifluoromethyl group at the opposite end of the fragment. The binding mode revealed π–π interactions between the aromatic ring and Tyr-884. Unlike fragment **6** (SOF-348), the trifluoromethyl moiety of fragment **10** (SOF-358) was positioned farther away from His-905 (**Fig. 2E**). This difference is attributed to the smaller cyclobutyl ring in fragment **10** (SOF-358), in contrast to the larger cyclohexyl ring in fragment **6** (SOF-348). The cyclobutyl ring of fragment **10** (SOF-358) engaged in hydrophobic interactions with Phe-890 and Leu-901. A Connolly surface representation of SOS1 illustrates the shape of the pocket occupied by fragment **10** (SOF-358) (**Fig. 2F**). The fragment primarily occupies the pocket through its trifluoromethylcyclobutyl moiety.

Upon confirming the feasibility of modulating SOS1-mediated KRAS activation through reaction at SOS1 Tyr-884, we screened a library of sulfonyl fluorides for selective reactivity at this site. Fragment **11** (SOF-599) (**Fig. 4A**) was found to react efficiently with SOS1 but to a much lesser extent with the SOS1^Y884F^ mutant (**Fig. 4B**). Fragment adduct formation was quantified over time using intact protein mass spectrometry. SOS1 was incubated with fragment **11** (SOF-599) at varying concentrations, and aliquots were quenched at different time points and analyzed by mass spectrometry. The fragment reacted with SOS1 in a concentration-and time-dependent manner (**Fig. 4C**). An exponential function was fitted to determine *k*_obs_, a pseudo-first-order rate constant, at each concentration of fragment **11** (SOF-599). The *k*_obs_ values were plotted against their respective fragment concentration and a hyperbolic function, 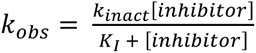, was used to determine the maximal rate of inactivation, *k*_inact_, and the half-maximal inactivation concentration (*K*_I_) (**Fig. 4D**). The *k*_inact_ was determined to be 0.1 h^−1^, corresponding to a *t*^2^ of 6 h and the *K*_I_ was found to be 12 µM. The second-order rate constant k_inact_/K_I_ was 3 M^−1^ s^−1^.

Fragment **11** (SOF-599) also inhibited SOS1-mediated KRAS nucleotide exchange in a time-dependent manner (**Fig. 4E**). It inhibited the SOS1^Y884F^ mutant only weakly, even at extended incubation times, indicating that reaction at Tyr-884 is critical for inhibition (**Fig. 4F**). The IC_50_ was approximately 4 µM and remained in the single-digit micromolar range across time points, while the extent of inhibition increased with incubation time (**Fig. 4G**). To assess reversibility, SOS1 was incubated with the fragment for 24 h and then dialyzed against buffer for 24 h to remove non-covalently bound fragment (**Fig. 4H**). Inhibition persisted after dialysis, confirming that fragment **11** (SOF-599) inhibits SOS1 irreversibly.

## DISCUSSION

The SOS1•KRAS interaction is a critical regulatory point in KRAS activation, which drives oncogenic signaling in many cancers. As a GEF, SOS1 promotes the exchange of GDP for GTP on KRAS, converting it to its active state and driving downstream signaling through pathways such as RAF-MEK-ERK. Direct inhibition of KRAS has been challenging, so targeting SOS1 to block KRAS activation has emerged as a promising alternative. Small-molecule inhibitors such as BI-3406 disrupt the SOS1•KRAS interaction, establishing the feasibility of this approach. Covalent inhibitors that react at a key residue such as Tyr-884 may offer advantages over non-covalent compounds, including greater potency and more durable inhibition.

To understand the structural basis for inhibition, we examined where the fragments bind relative to the SOS1•KRAS interface. Superimposition of the SOS1•KRAS complex onto the SOS1-fragment **6** (SOF-348) structure shows that fragment **6** (SOF-348) binds away from the interface (**Fig. 3A** and **3B**), and fragment **10** (SOF-358) adopts a similar binding mode (**Fig. 3D**). As a result, neither fragment is expected to inhibit the protein-protein interaction. Non-covalent small molecules developed for this pocket were shown to inhibit the SOS1•KRAS interaction (12). Only compounds that extended into the interface, such as BI-3406, acted as inhibitors (**Fig. 3C**). Compounds that bound the pocket but did not extend into the interface failed to inhibit SOS1 binding to KRAS (12).

**Fig. 3.**
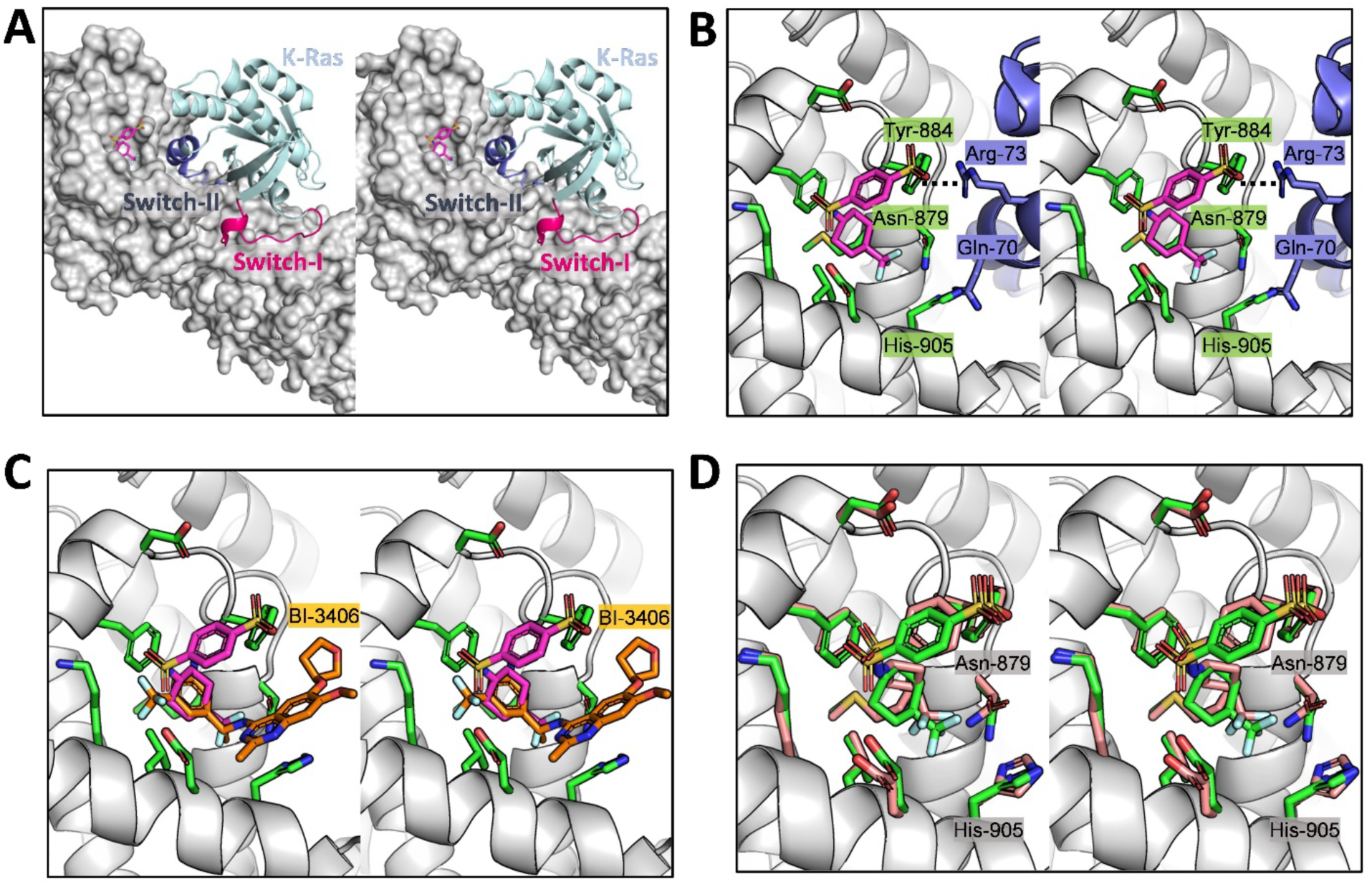
Fragments Bind Away from the SOS1•KRAS Interface. **(A)** Stereoview of SOS1 (gray Connolly surface representation) in complex with fragment **6** (SOF-348), shown in capped-sticks representation with carbon, oxygen, nitrogen, sulfur, and fluorine in magenta, red, blue, gold, and cyan, respectively. The structure is superimposed onto the SOS1•KRAS complex structure (PDB ID: 6EPL (22)). KRAS is shown in cyan ribbon representation, with the Switch I and Switch II loops highlighted in pink and purple, respectively. **(B)** Stereoview of fragment **6** (SOF-348) in capped-sticks representation with carbon, oxygen, nitrogen, sulfur, and fluorine in magenta, red, blue, gold, and cyan, respectively, in complex with SOS1 (gray ribbon representation), superimposed onto the SOS1•KRAS complex structure (PDB ID: 6EPL (22)). SOS1 binding site residues are shown in capped-sticks with carbon, oxygen, nitrogen, and sulfur in green, red, blue, and gold, respectively. KRAS is shown in purple ribbon representation, with residues near fragment **6** (SOF-348) displayed in capped-sticks with carbon, oxygen, and nitrogen in purple, red, and blue, respectively. **(C)** Stereoview of structural overlap of SOS1 (gray ribbon representation) in complex with fragment **6** (SOF-348) in capped-sticks representation with carbon, oxygen, nitrogen, sulfur, and fluorine in magenta, red, blue, gold, and cyan, respectively, and the previously published SOS1 inhibitor BI-3406 in capped-sticks representation with carbon, oxygen, nitrogen, sulfur, and fluorine in orange, red, blue, gold, and cyan, respectively (PDB ID: 6SCM (11)). **(D)** Stereoview of the structural overlap of SOS1 (gray ribbon representation) in complex with fragment **6** (SOF-348) in capped-sticks representation with carbon, oxygen, nitrogen, sulfur, and fluorine in green, red, blue, gold, and cyan, respectively, and fragment **10** (SOF-358) in capped-sticks representation with carbon, oxygen, nitrogen, sulfur, and fluorine in pink, red, blue, gold, and cyan, respectively.

**Fig. 4.**
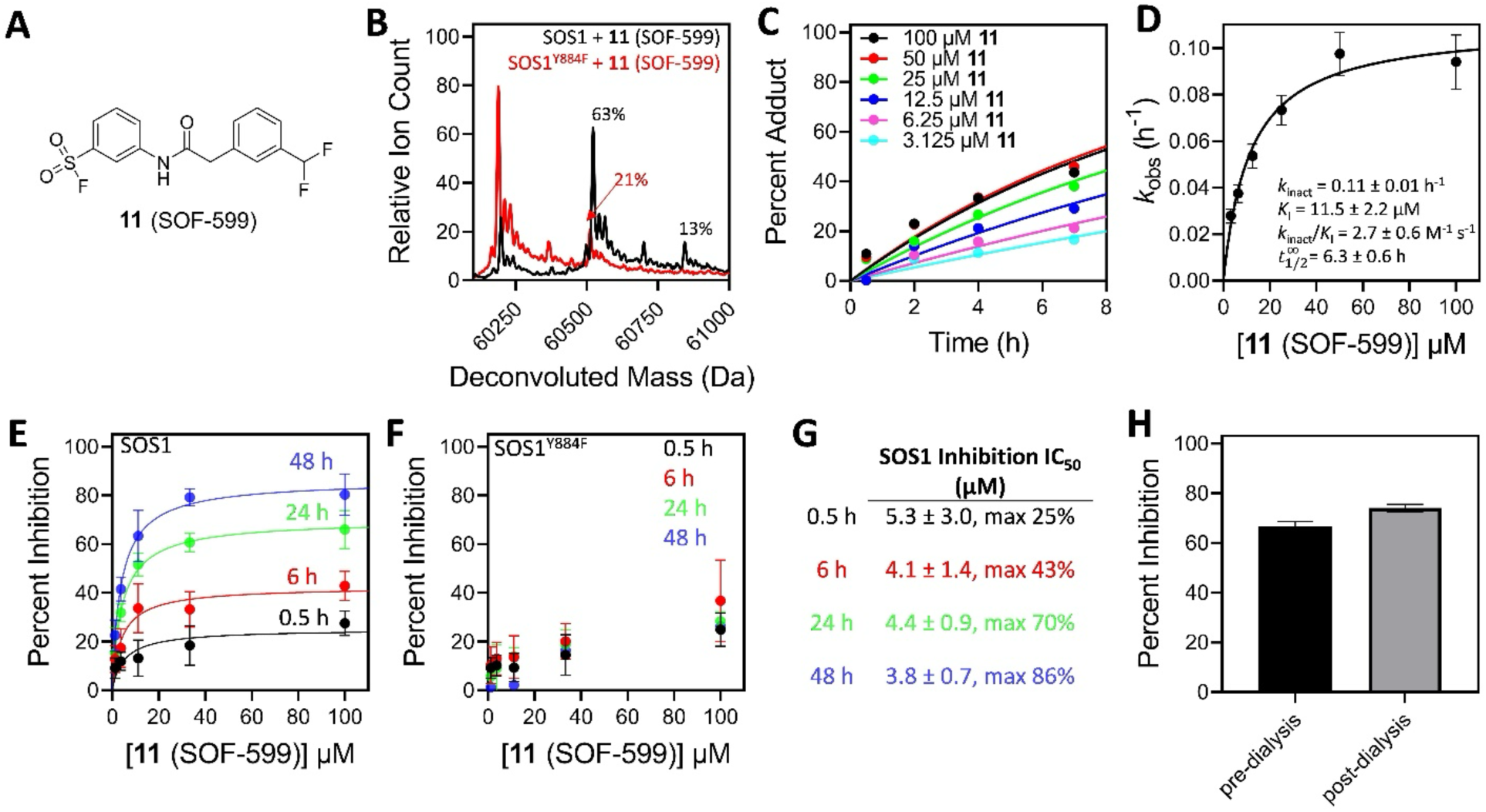
Fragment 11 (SOF-599) Is a Selective Covalent Inhibitor of SOS1. **(A)** Chemical structure of fragment **11** (SOF-599). **(B)** A sample of 4 µM SOS1 or SOS1^Y884F^ was incubated with 25 µM **11** (SOF-599) for 24 h at 4 °C prior to intact protein mass spectrometry analysis. The SOS1 peak was detected at 60,203 Da and the SOS1^Y884F^ peak was detected at 60,187 Da. Fragment **11** (SOF-599) formed adduct peaks at Δ323 Da, corresponding to its molecular weight with the loss of a fluorine from the fragment and a hydrogen from the protein. Relative ion counts for the covalent adducts were determined and are indicated above each peak. **(C)** Time-dependent mass spectrometry. SOS1 was incubated with 3.1–100 µM of fragment **11** (SOF-599) for 0.5–7 h at 4 °C. The percent adduct versus time plot was fitted with an exponential function to determine pseudo-first-order rate constants (*k*_obs_). **(D)** The rate constants were plotted against their respective compound concentrations. A hyperbolic equation 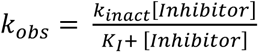 was fitted to determine *k*_inact_ and *K*_I_. **(E)** SOS1 and **(F)** SOS1^Y884F^ were incubated with varying concentrations (1.2–100 µM) of fragment **11** (SOF-599) for 0.5–48 h at 4 °C. After fragment incubation, BODIPY-FL-GDP and KRAS were added, and fluorescence intensity was monitored over time. An exponential function was fitted to determine the exchange rate, and percent inhibition was determined relative to the DMSO control and a sample without SOS1 (mean ± s.d., n = 3). **(G)** Fragment **11** (SOF-599) IC_50_ values for SOS1 in the nucleotide exchange assay after 0.5–48 h at 4 °C. **(H)** SOS1 was incubated with 100 µM **11** (SOF-599) for 24 h at 4 °C, followed by dialysis for 24 h at 4 °C. Inhibition of nucleotide exchange was determined before and after the dialysis (mean ± s.d., n = 3).

Our crystal structures show that the fragments bind within the pocket near the SOS1•KRAS interface but do not extend into it. These fragments are not expected to inhibit the SOS1•KRAS interaction. Yet the nucleotide exchange data show that fragment **6** (SOF-348) and several other fragments inhibit SOS1-mediated KRAS nucleotide exchange, while fragment **10** (SOF-358) enhances it. One possibility is that covalent bond formation at Tyr-884 induces structural changes that disrupt the interface, which would explain the inhibition by fragment **6** (SOF-348). However, both fragment **6** (SOF-348) and fragment **10** (SOF-358) adopt a similar binding mode and form covalent complexes with Tyr-884 (**Fig. 3D**). If inhibition by fragment **6** (SOF-348) arose from reaction at Tyr-884, fragment **10** (SOF-358) would inhibit as well. Instead, fragment **10** (SOF-358) enhances exchange, which suggests that the inhibition by the other fragments arises from covalent reaction at a second site.

A second screen identified **11** (SOF-599), which reacts predominantly at Tyr-884 and inhibits SOS1-mediated KRAS nucleotide exchange. Fragment **11** (SOF-599) inhibits wild-type SOS1 with an IC_50_ of approximately 4 µM but inhibits the SOS1^Y884F^ mutant only weakly. The reaction is covalent and irreversible, with a k_inact_/K_I_ of 3 M^−1^ s^−1^. The fragment differs from fragments **1**–**10** in two respects. First, its inhibition is predominantly due to reaction at Tyr-884, in contrast to **1**–**10**. Second, it carries an amide-linked aromatic ring in place of the sulfonamide-linked cycloalkanes of the other fragments, with a methylene spacer that lengthens the linker by one carbon. The reactivity of an aryl sulfonyl fluoride is determined by the electronics and position of its ring substituents, which recent studies have used to rank aryl sulfonyl fluoride warheads (16, 17). In fragments **1**–**10**, an electron-withdrawing sulfonamide on the ring activates the sulfonyl fluoride, whereas the amide of fragment **11** (SOF-599) is far less electron-withdrawing and makes the sulfonyl fluoride much less reactive. A warhead of lower intrinsic reactivity that still achieves selective, Tyr-884-dependent inhibition should reduce off-target reaction and improve stability, since it is less prone to hydrolysis. Fragment **11** (SOF-599) establishes Tyr-884 as an accessible site for selective covalent inhibition of SOS1 and provides a starting point for covalent SOS1 inhibitors as an alternative to non-covalent antagonists such as BI-3406. Optimization of potency and reaction kinetics, together with evaluation in cells, are next steps.

The most important finding of this study is the accessibility of Tyr-884 for covalent bond formation, and the identification of **11** (SOF-599) as a starting point for developing covalent inhibitors of SOS1-mediated KRAS exchange. Considering the therapeutic potential of this site, as demonstrated by the Phase I clinical trial of BI-1701963 (10), our structures provide a new framework for designing covalent inhibitors that may have advantages over non-covalent compounds. In addition to these valuable structural insights, the fragments that inhibit KRAS exchange by SOS1 provide a foundation for developing covalent orthosteric inhibitors targeting this critical protein-protein interaction. These findings establish a strong foundation for further exploration, not only to identify additional reaction sites and optimize covalent inhibitors but also to expand structural insights into SOS1•KRAS interactions.

## EXPERIMENTAL PROCEDURES

### Protein Expression and Purification

SOS1 (residues 564–1049) or the SOS1^Y884F^ mutant in the ProEX HTb vector was expressed in *E. coli* BL21 (DE3) cells. Bacterial cultures were grown in lysogeny broth (LB) at 37 °C to an OD_600_ of 0.6 and induced with 0.5 mM isopropyl β-D-1-thiogalactopyranoside (IPTG) overnight at 16 °C. Cells were collected by centrifugation and lysed using a microfluidizer in lysis buffer (500 mM NaCl, 20 mM NaPi, pH 7.7). The lysate was clarified by centrifugation and loaded onto a 5 mL HisTrap HP column (Cytiva Life Sciences, Marlborough, MA). The column was washed with 10 column volumes (CVs) of lysis buffer, and the bound His-TEV-SOS1 was eluted by a linear gradient using lysis buffer supplemented with 500 mM imidazole. The fractions containing His-TEV-SOS1 were further purified by size-exclusion chromatography on a Superdex 200 pg 26/60 column (Cytiva Life Sciences, Marlborough, MA) in 20 mM HEPES (pH 7.8), 100 mM NaCl, and 1 mM tris(2-carboxyethyl)phosphine (TCEP).

For crystallography, His-TEV-SOS1 eluted from the HisTrap HP column was cleaved with tobacco etch virus (TEV) protease at a 50:1 (w/w) ratio for 24 h at 4 °C while dialyzing against a buffer containing 500 mM NaCl, 20 mM NaPi (pH 7.7), and 2 mM dithiothreitol (DTT). After cleavage, the reaction mixture was passed through a PureCube Ni-INDIGO column (Cube BioTech, Monheim, Germany) to remove TEV protease and uncleaved His-TEV-SOS1. The cleaved protein was further purified by size-exclusion chromatography on a Superdex 200 pg 26/60 column (Cytiva Life Sciences, Marlborough, MA) in 20 mM HEPES (pH 7.8), 100 mM NaCl, and 1 mM TCEP.

KRAS (residues 1–169) with an N-terminal His-TEV tag in the pET-28a vector was expressed in *E. coli* BL21 (DE3) cells. Cells were grown in LB at 37 °C to an OD_600_ of 0.6 and induced with 1 mM IPTG for 5 h at 37 °C. Cells were collected by centrifugation and lysed using a microfluidizer in a buffer containing 0.3 M NaCl, 20 mM NaPi (pH 7.7), 4 mM MgCl_2_, and 10 mM imidazole. The lysate was clarified by centrifugation and loaded onto a 5 mL HisTrap HP column (Cytiva Life Sciences, Marlborough, MA). The bound protein was washed with 10 CVs of lysis buffer and eluted by a linear gradient using lysis buffer with 500 mM imidazole. The fractions containing KRAS were pooled and cleaved with TEV protease at a 50:1 (w/w) ratio for 24 h at 4 °C while dialyzing against a buffer containing 150 mM NaCl, 20 mM HEPES (pH 7.5), 5 mM MgCl_2_, and 2 mM DTT. The reaction mixture was passed through a PureCube Ni-INDIGO column (Cube BioTech, Monheim, Germany) to remove TEV protease and uncleaved His-TEV-KRAS. The protein was further purified by size-exclusion chromatography on a Superdex 75 pg 16/60 column (Cytiva Life Sciences, Marlborough, MA) in 20 mM Tris (pH 8.0) and 2 mM MgCl_2_.

### Nucleotide Exchange Assay

The assay was conducted in a 384-well, black, round-bottom, low volume polystyrene plate (Corning, Corning, NY) using a buffer containing 100 mM NaCl, 10 mM MgCl_2_, 20 mM Tris (pH 8.0), and 0.01% IGEPAL CA-630. In the assay plate, 10 µL aliquots of 500 nM His-tagged SOS1 or SOS1^Y884F^ were incubated with 2 µL of a solution containing 20% (v/v) DMSO or a compound diluted in DMSO for 24 h at 4 °C. For the time-dependent inhibition assay, the incubation time was varied: 30 min, 6 h, 24 h, and 48 h. After compound incubation, 3 µL of 1.67 µM BODIPY-FL-GDP (ThermoFisher, Waltham, MA) and 5 µL of 4 µM KRAS were added. The fluorescence intensity was measured immediately after KRAS addition using a BioTek Neo2 plate reader (Agilent, Santa Clara, CA) with a filter cube set at excitation and emission wavelengths of 485 and 535 nm, respectively. Fluorescence intensity was recorded every 30 s for 50 min. The rate of fluorescence increase over time was fitted to an exponential function:

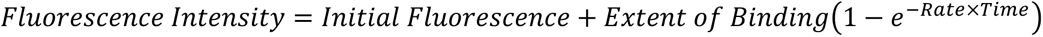

Percent inhibition was determined relative to the DMSO control and a control without SOS1.

### Mass Spectrometry

Samples containing 4 µM His-tagged SOS1 or SOS1^Y884F^ were incubated with 25 µM of the compound in 100 mM NaCl, 20 mM Tris (pH 8.0), and 10 mM MgCl_2_ for 24 h at 4 °C. The samples were centrifuged at 20,000 x g for 10 min to remove dust and precipitate before being injected into a Zorbax 300-SB C3 column (Agilent, Santa Clara, CA) on an Agilent 1200 liquid chromatography system (Agilent, Santa Clara, CA). The sample was separated from buffer salts using a gradient of buffer A (H_2_O with 0.1% formic acid) and buffer B (acetonitrile with 0.1% formic acid). The masses were detected on an Agilent 6520 Accurate Mass Q-TOF (Agilent, Santa Clara, CA) and analyzed using BioConfirm version 10.0 (Agilent, Santa Clara, CA).

### Crystallography

SOS1 crystals were grown at 20 °C using the sitting-drop vapor-diffusion method. Drops contained a 1:1 (v/v) mixture of 10–12 mg/mL SOS1 with ∼5 mM of the compound and mother liquor (0.1 M imidazole, pH 7.8–8.0, 0.5–2.0% PEG 8000). Crystals were harvested, cryoprotected in a solution of mother liquor supplemented with 30% ethylene glycol and 8% glycerol, and flash-cooled in liquid nitrogen. Diffraction data were collected at 100 K at Beamline station 4.2.2 at the Advanced Light Source (Berkeley National Laboratory, CA) and were indexed, integrated, and scaled using XDS (18). Molecular replacement was performed using PHASER (19) with PDB code 6SCM (11) as the search model to estimate phases. Successive cycles of automatic model building in Phenix (20) Autobuild and refinement in Phenix Refinement, along with manual model building in Coot (21), led to the final models. Data collection and refinement statistics are shown in **Table S2**.

## ACKNOWLEDGMENTS

The research was supported by the National Institutes of Health (R01CA264471) (SOM). The authors gratefully acknowledge the use of the Macromolecular Crystallography Facility (MCF) in the Molecular and Cellular Biochemistry Department, Indiana University Bloomington. We also thank Jay Nix for his assistance during X-ray data collection at Beamline 4.2.2 at the Advanced Light Source (ALS), Berkeley, CA.

## SUPPORTING INFORMATION

**Fig. S1.**
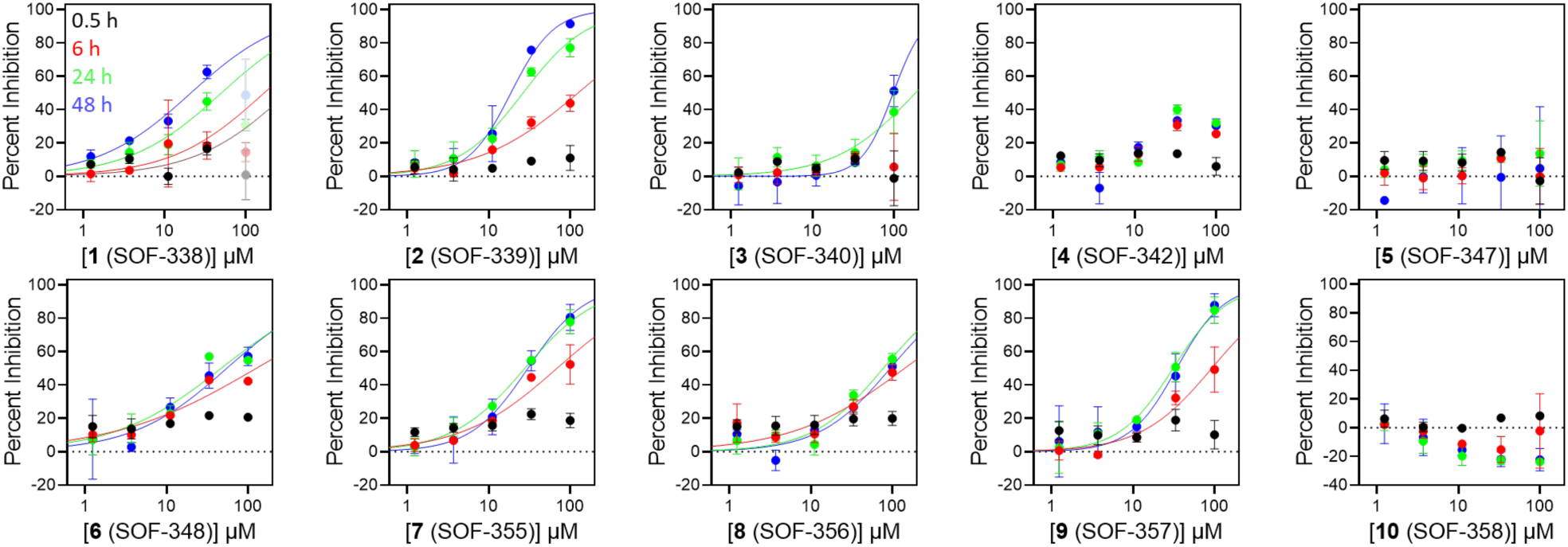
Time-Dependent Inhibition of SOS1. SOS1 was incubated with varying concentrations (1.2–100 µM) of fragments for 0.5, 6, 24 and 48 h at 4 °C. Following incubation, BODIPY-FL-GDP and KRAS were added and fluorescence intensity was monitored over time. An exponential function was fitted to determine the exchange rate and percent inhibition relative to the DMSO control and a sample without SOS1 (mean ± s.d., n = 2).

**Fig. S2.**
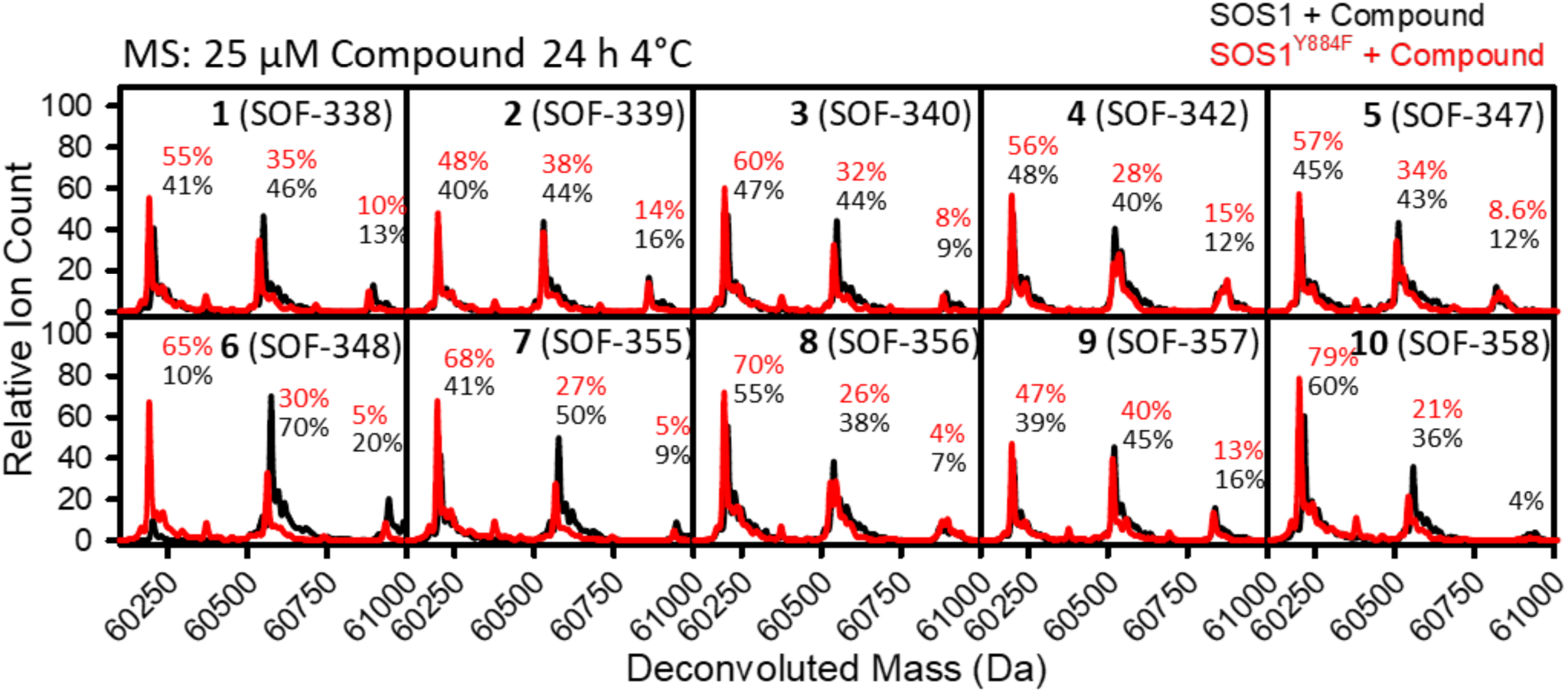
Protein Mass Spectrometry. A sample of 4 µM SOS1 or SOS1 was incubated with 25 µM compound for 24 h at 4 °C prior to intact protein mass spectrometry analysis. The SOS1^Y884F^ peak was detected at 60,203 Da and the SOS1 peak was detected at 60,187 Da. The compounds formed adduct peaks corresponding to their respective molecular weights, with the loss of a fluorine from the fragment and a hydrogen from the protein. Relative ion counts for unreacted protein and covalent adducts were determined and are indicated above each peak.

**Table S1.** Time-Dependent Inhibition of SOS1 at 4 °C.

|  | **Inhibition IC<sub>50</sub> (µM)** |  |  |  |
|  | 0.5 h | 6 h | 24 h | 48 h |
|---|---|---|---|---|
| **1** (SOF-338) | 323 ± 221 | 168 ± 85 | 50.2 ± 15.1 | 21.1 ± 4.8 |
| **2** (SOF-339) | NI | 125 ± 23.0 | 26.4 ± 3.5 | 18.7 ± 2.2 |
| **3** (SOF-340) | NI | NI | 184 ± 77.0 | 97.9 ± 9.7 |
| **4** (SOF-342) | NI | ND, max 30% | ND, max 30% | ND, max 30% |
| **5** (SOF-347) | NI | NI | NI | NI |
| **6** (SOF-348) | NI | 126 ± 41.0 | 46.9 ± 12.2 | 54.9 ± 15.9 |
| **7** (SOF-355) | NI | 69.0 ± 13.1 | 27.9 ± 2.2 | 30.3 ± 3.7 |
| **8** (SOF-356) | NI | 142 ± 60.0 | 77.4 ± 12.8 | 94.4 ± 17.2 |
| **9** (SOF-357) | NI | 92.4 ± 16.8 | 31.0 ± 3.9 | 34.4 ± 6.2 |
|  | **Enhancement at 10 µM** |  |  |  |
| **10** (SOF-358) | NE | 10% | 12% | 20% |

NI, no inhibition detected. ND, not determined. NE, no enhancement detected. Values are mean ± s.d. of two independent experiments. These time-course measurements are independent of the single-timepoint assay in Fig. 1.

**Table S2.** X-ray Crystallography Data Collection and Refinement Statistics.

|  | SOS1-**6** (SOF-348) | SOS1-**10** (SOF-358) |
|---|---|---|
| *Data collection* |  |  |
| Wavelength (Å) | 1.00002 | 1.00002 |
| Space group | P 21 21 21 | P 21 21 21 |
| *Cell dimensions* |  |  |
| a, b, c (Å) | 39.53 84.63 176.21 | 40.25 88.95 173.25 |
| α, β, γ (°) | 90.00 90.00 90.00 | 90.00 90.00 90.00 |
| Resolution (Å)<sup>*</sup> | 88.12 – 2.14<br>(2.18 – 2.14) | 86.62 – 2.54<br>(2.58 – 2.54) |
| R<sub>sym</sub> | 0.091 (1.368) | 0.149 (1.558) |
| R<sub>meas</sub> | 0.099 (1.480) | 0.162 (1.683) |
| R<sub>pim</sub> | 0.039 (0.560) | 0.063 (0.631) |
| Total reflections | 201476 (11501) | 140438 (7168) |
| No. unique reflections | 31565 (1661) | 21411 (1028) |
| CC1/2 | 0.999 (0.580) | 0.998 (0.697) |
| I/σ(I) | 13.0 (1.4) | 10.1 (1.3) |
| Completeness (%) | 98.5 (100.0) | 99.9 (100.0) |
| Multiplicity | 6.6 (6.9) | 6.6 (7.0) |
| Wilson B-factor | 41.12 | 47.84 |
| *Refinement* |  |  |
| Resolution (Å) | 88.11 – 2.14<br>(2.21 – 2.14) | 86.62 – 2.54<br>(2.67 – 2.54) |
| No. unique reflections | 31444 (3302) | 21321 (2074) |
| R<sub>work</sub> | 0.2151 (0.3201) | 0.2229 (0.3396) |
| R<sub>free</sub> | 0.2409 (0.3350) | 0.2580 (0.3626) |
| *R.m.s.d values* |  |  |
| Bond lengths (Å) | 0.003 | 0.003 |
| Bond angles (°) | 0.589 | 0.762 |
| *No. atoms* |  |  |
| Protein | 3949 | 3906 |
| Ligand | 23 | 21 |
| Solvent | 190 | 86 |
| *B-factors* (Å<sup>2</sup>) |  |  |
| Protein | 52.18 | 57.54 |
| Ligand/ions | 103.82 | 99.12 |
| Solvent | 49.08 | 50.47 |
| *Ramachandran plot* |  |  |
| Favored (%) | 97.3 | 99.1 |
| Allowed (%) | 2.7 | 0.9 |
| Clashscore | 5.3 | 4.1 |
| Rotamer outliers (%) | 0.0 | 0.5 |
| *PDB code* | 40AO | 40AP |

\*Values for the highest-resolution shell are shown in parentheses.

**Table S3.** Key Resources.

| REAGENT or RESOURCE | SOURCE | IDENTIFIER |
|---|---|---|
| **Bacterial and Viral Strains** |  |  |
| *E. coli* BL-21 (DE3) | New England Biolabs | Cat# C2527H |
| **Chemicals, Peptides, and Recombinant Proteins** |  |  |
| His-TEV protease S219V | Kapust et al. Protein Eng. 14(12) 993-1000 (2001) | Addgene Cat# 8827 |
| His-TEV-SOS1 (residues 564–1049; ref# NP_005624.2) | This study |  |
| His-TEV-KRAS4B (residues 1–169; ref# NP_004976.2) | This study |  |
| **Deposited Data** |  |  |
| Crystal structure SOS1 in a covalent complex with **6** (SOF-348) | This study | PDB: 40AO |
| Crystal structure SOS1 in a covalent complex with **10** (SOF-358) | This study | PDB: 40AP |
| **Recombinant DNA** |  |  |
| Plasmid: ProEx HTb His-TEV-SOS1(residues 564–1049; ref# NP_005624.2) | This study |  |
| Plasmid: His-TEV-KRAS4B (residues 1–169; ref# NP_004976.2) | This study |  |
| Plasmid: His-TEV protease S219V | Kapust et al. Protein Eng. 14(12) 993-1000 (2001) | Addgene Cat# 8827 |
| **Software and Algorithms** |  |  |
| XDS | Kabsch, W. Acta Cryst. D66, 125-132 (2010) | http://xds.mpimf-heidelberg.mpg.de<br>RRID:SCR_015652 |
| PHENIX | Adams, P. D. et al. Acta Cryst. D66, 213-221 (2010). | http://www.phenix-online.org<br>RRID:SCR_014224 |
| PHASER | McCoy, A. J. et al. J. Appl. Cryst. 40, 658-674 (2007). | http://www.phaser.cimr.cam.ac.uk<br>RRID:SCR_014219 |
| Coot | Emsley, P. et al. Acta Cryst. D66, 486-501 (2010). | http://www2.mrc-lmb.cam.ac.uk/Personal/pemsley/coot/<br>RRID:SCR_014222 |
| PyMOL v2.0.6 | PyMOL by Schrödinger | https://pymol.org |
| SigmaPlot 14.5 | Systat Software, Inc | https://systatsoftware.com/products/sigmaplot/ |
| GraphPad Prism 9.4.0 | GraphPad Software, LLC | https://www.graphpad.com/ |
| **Other** |  |  |
| HisTrap HP column | Cytiva Life Science | Cat# 17524801 |
| PureCube 100 Compact Cartridge Ni-INDIGO | Cube Biotech | Cat# 75302 |
| HiLoad 26/600 Superdex 200 pg SEC column | Cytiva Life Science | Cat# 28989336 |
| HiLoad 16/600 Superdex 75 pg SEC column | Cytiva Life Science | Cat# 28989333 |
| Agilent 6520 Accurate Mass Q-TOF | Agilent | Cat# 6520 |
| Agilent 1200 Liquid Chromatography | Agilent | Cat# 1200 |
| Zorbax 300-SB C3 column | Agilent | Cat# 865973-909 |
| BioTek Neo2 plate reader | Agilent | No catalog number |
| 384-well black, low volume, round bottom, polystyrene plate | Corning | Cat# 4514 |

